# Improved prediction of femoral fracture toughness in mice by combining standard medical imaging with Raman spectroscopy

**DOI:** 10.1101/2020.11.20.391383

**Authors:** Christine Massie, Emma Knapp, Keren Chen, Andrew J Berger, Hani A Awad

## Abstract

Bone fragility and fracture risk are assessed by measuring the areal bone mineral density (aBMD) using dual-energy X-ray absorptiometry (DXA). While aBMD correlates with bone strength, it is a poor predictor of fragility fracture risk. Alternatively, fracture toughness assesses the bone’s resistance to crack propagation and fracture, making it a suitable bone quality metric. Here, we explored how femoral midshaft measurements from DXA, micro-computed tomography (μCT), and Raman spectroscopy could predict fracture toughness. We hypothesized that ovariectomy (OVX) decreases aBMD and fracture toughness compared to controls and we can optimize a multivariate assessment of bone quality by combining results from X-ray and Raman spectroscopy. Female mice underwent an OVX (n=5) or sham (n=5) surgery at 3 months of age. Femurs were excised 3 months after ovariectomy and assessed with Raman spectroscopy, μCT, and DXA. Subsequently, a notch was created on the anterior side of the mid-diaphysis of the femurs. Three-point bending induced a controlled fracture that initiated at the notch. The OVX mice had a significantly lower aBMD, cortical thickness, and fracture toughness when compared to controls (p<0.05). A leave one out cross-validated (LOOCV) partial least squares regression (PLSR) model based only on the combination of aBMD and cortical thickness showed no significant predictive correlations with fracture toughness, whereas a PLSR model based on principal components derived from the full Raman spectra yielded significant prediction (r^2^=0.71, p<0.05). Further, the PLSR model was improved by incorporating aBMD, cortical thickness, and principal components from Raman spectra (r^2^=0.92, p<0.001). This exploratory study demonstrates combining X-ray with Raman spectroscopy leads to a more accurate assessment of bone fracture toughness and could be a useful diagnostic tool for the assessment of fragility fracture risk.

## Introduction

Osteoporosis is a global epidemic with a socioeconomic burden that escalates at an alarming rate with the increase in life expectancy of the aging population (Wright et al., 2014). The risk of this disease doubles every decade of life, such that a quarter of women over 65 years of age are osteoporotic (Force et al., 2018). The clinical consequences of this disease, which is characterized by bone loss and low bone mineral density (BMD), include fragility fractures that lead to morbidity and mortality. Unfortunately, the bone mass-based screening of osteoporosis and osteopenia often does not accurately identify patients at high risk before a fragility fracture occurs. The underlying issue leading to these fractures is the deteriorating mechanical properties, which is not detected by BMD. While BMD has been shown to correlate sufficiently well with bone strength and modulus (Choi et al., 1990; Keaveny et al., 1994), it is not a reliable independent determinant of fracture toughness (Inzana et al., 2013) whether measured by dual-energy X-ray absorptiometry (DXA) or other imaging modalities such as quantitative computed tomography (Wang et al., 1998) or fracture risk (McClung, 2005). Opposed to BMD, fracture risk is more dependent on mechanistic pathways such as fracture toughness, strength, or fatigue strength (Hernandez and Keaveny, 2006).

There is a critical need for additional assessments to complement BMD and enhance diagnostic sensitivity. Alternatively, X-ray computed tomography (CT) is a high-resolution imaging modality which analyzes 3D bone structures. In efforts to increase sensitivity for fracture risk, CT scans have been coupled with finite element analysis to computationally derive biomechanical properties. However, this approach has been shown to have comparable sensitivity and specificity to DXA (Adams et al., 2018) for identifying high fracture risk patients. As an alternative to medical imaging, Raman spectroscopy is a non-destructive vibrational spectroscopy technique that can reveal important compositional information about bone matrix and is sensitive to chemical and structural changes in the matrix (Morris and Mandair, 2011). Raman spectroscopy has been used to study the chemical composition of bones (Carden and Morris, 2000), the effect of glucocorticoid in rheumatoid arthritis (Maher et al., 2011), aging bone (Ager et al., 2005) and healing (Ahmed et al., 2018), but has yet to be used clinically. Therefore, noninvasive metrics for bone quality and fragility fracture risk based on imaging alone remain elusive.

While not clinically feasible, measuring the tissue’s mechanical properties would improve bone quality assessment in experimental preclinical models. These include strength and elasticity, which are typically measured in human cadaver specimens and animal models by loading the bone to failure in bending, torsion, or compression (Cole and van der Meulen, 2011). These properties, however, do not capture the bone’s ability to dissipate high stress concentrations associated with internal microscopic cracks that initiate and progressively propagate during daily cycles of subfailure fatigue loading. Alternatively, fracture toughness tests determine the ability of the tissue to resist brittle fractures arising from the propagation of an existing flaw under loading. Fracture toughness generally does not correlate with the bone’s standard mechanical properties (strength and elasticity) (Ritchie, 2011), but it is arguably more reflective of bone quality and predictive of fragility fracture risk (Launey et al., 2010; Ritchie, 2010; Ritchie, 2011). We have previously adapted a fracture toughness model from Ritchie et al. (2008) (Ritchie et al., 2008) and demonstrated that Raman spectroscopy was predictive of fracture toughness (Inzana et al., 2013). While such correlations are intriguing, the diagnostic utility of Raman spectroscopy depends on its ability to classify bone health according to their fracture toughness.

In this study, we hypothesized that in mice ovariectomy (OVX) induces bone loss leading to decreased fracture toughness compared to sham controls, and a multivariate assessment of bone quality including X-ray and Raman would accurately predict OVX-associated changes in fracture toughness. OVX is a well-defined model for postmenopausal osteoporosis. Several studies have shown that OVX leads to a decrease in bone qualities such as bone volume fraction (Bouxsein et al., 2005), femoral BMD measured by DXA (Lei et al., 2009), bone volume of the distal metaphysis of trabecular bone (Rosales Rocabado et al., 2018), and trabecular thickness (Anbinder et al., 2016). Interestingly, the cortical bone response to OVX is not well defined (Osterhoff et al., 2016). However, fractures not involving vertebral bodies are predominantly cortical (Zebaze et al., 2010) and are considered to be a leading cause of morbidity and mortality in the osteoporotic elderly (Zebaze et al., 2010). Furthermore, there is a gap in assessing the effects of OVX-induced bone loss on fracture toughness, which we attempt to address by creating a model to predict such changes with X-ray and Raman.

## Methods

### Animals and Surgical Procedures

Animal studies were performed with adherence to protocols approved by the University of Rochester’s Committee on Animal Resources. Three-month-old C57Bl/6J female mice were sedated using a ketamine/xylazine (60-100 mg/kg and 10 mg/kg body weight dosage, respectively) and underwent either an ovariectomy or a sham surgery. Briefly, a small incision was made on the dorsal side of the mice, proximal of the hip joints. Ovary fat pads were located through the incision, and then placed back (for the sham group, n=5) or removed from the fat pad by cauterization (for the OVX group, n=5). The animals were sacrificed three months later, and the left and right femurs were collected for subsequent imaging, Raman spectroscopy, and biomechanical testing and averaged per mouse. The femurs were frozen at −20°C prior to and after testing.

### Raman Spectroscopy

As described previously (Maher and Berger, 2010; Maher et al., 2011), Raman measurements were performed *ex vivo* on the excised, thawed femurs using an 830 nm, 150 mW laser that was focused in a spot with 230-μm diameter. Three locations were measured and averaged along the anterior mid-diaphysis over a 3.5 mm^2^ area at 1 mm intervals to coregister with where the crack would be initiated for fracture toughness testing. The bone orientation was kept consistent for all measurements to control collagen fiber orientation. Excitation and detection were performed in a non-contact manner. For each location, spectra were acquired for five 60-second exposures. Spectral pre-processing involved cosmic ray removal, readout and dark current subtraction, and image aberration correction. The spectra (744-1740 cm^−1^) were fit to a 5^th^ order polynomial to remove fluorescence (Figure S1E). Spectra were smoothed with a Savitzky-Golay filter (Savitzky and Golay, 1964) over a window of 3 pixels (5cm^−1^). Spectra were normalized to their mean absolute deviation (MAD) with respect to their mean, determined by:

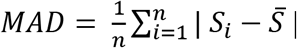

where S is the spectral intensity (Inzana et al., 2013) (Figure S1).

### Micro-Computed Tomography

The femurs were scanned by micro computed tomography (μCT; VivaCT 40; Scanco Medical; Bassersdorf, Switzerland) at a 10.5-micron isotropic resolution using an integration time of 300 ms, energy of 55 kVp, and intensity of 145 μA. The axial length of scans was 304.5 μm. The Scanco midshaft evaluation was used to determine the cortical thickness, volumetric bone mineral density (vBMD) and bone volume by averaging over 300 microns at the femoral midshaft. The cortical bone threshold was 330 which yields a linear attenuation coefficient of 2.64 and 582 mgHA/ccm and for trabecular bone threshold was 260 which yields a linear attenuation of 2.08 and 416 mgHA/ccm. 12 femurs were imaged simultaneously in special holders in air. All femurs were imaged in the same session in which a calibration was performed by scanning a phantom with 5 cylinders of different densities at 55 kV, 145 μA, and 250 ms integration time.

### Dual Energy X-ray Absorptiometry (DXA)

DXA measurements were taken using a Lunar PIXImus system (GE Lunar Corp; Madison, Wisconsin). Areal bone mineral density (aBMD) measurements were taken on the midshaft of the *ex vivo* femurs.

### Fracture Toughness

Methods previously published by our group (Inzana et al., 2013) were used to assess femoral fracture toughness. A razor blade was used to create a through-wall notch on the anterior surface of the femur, approximately at the mid-diaphysis. To create a smooth, polished notch, a 1μm diamond suspension was used. The notch depth was confirmed using X-ray (UltraFocus; Faxitron; Tucson, Arizona). After the notch was created, the femurs were stored at room temperature for 2 hours while submerged in PBS. The samples underwent three-point bending tests, using an ElectroPuls E10000 (Instron; Norwood Massachusetts). To align the top bending post with the notch, a stereomicroscope was used. The testing set-up and femurs were submerged in PBS throughout the tests, with the notched, anterior side in tension. Testing was performed using a 6mm span and a displacement rate of 0.001mm/s along the posterior-anterior direction.

Following testing, the femurs were fixed, dehydrated, and gold sputter coated for scanning electron micrograph imaging (SEM; LEO 982 FE-SEM; Carl Zeiss SMT; Thornwood, NY). The SEM imaged the cross-section of the fractured femurs in the plane of the notch to allow for measurements of the half-crack angle at fracture instability, average cortical thickness, and mean radii. These measurements were taken on the SEM images using a custom created MATLAB^®^ code (R2016a; MathWorks Inc.; Natick, MA). The fracture toughness of each sample was then calculated using the method adapted from Ritchie et al. (Ritchie et al., 2008):

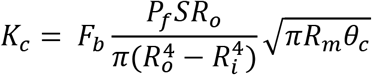

where *P*_*f*_ is the load at fracture, *S* is the span length, *R*_*o*_ and *R*_*i*_ are the mean outer and inner radii, *R*_*m*_ is the average of *R*_*o*_ and *R*_*i*_, *θ*_*C*_ is the half-crack angle at fracture instability, and *F*_*b*_ is a geometry factor for a thick-walled pipe as defined by Takahashi (Takahashi, 2002).

### Analysis

All statistical analyses were performed in MATLAB^®^ or JMP®14 (Version 14, SAS Institute, Cary, North Carolina). To compare the average effects of ovariectomy on experimentally measured parameters, a permutation test was used since it is a robust statistical test despite a small sample size and requires fewer assumptions than parametric tests (Collingridge, 2013; Gadbury et al., 2003; Önder et al., 2003; Routledge, 1997; Wilcox, 2003). Briefly, permutation testing works by randomly exchanging labels on experimentally observed data points from two treatment groups when performing significance tests and tests the null hypothesis that the treatment groups or labels are exchangeable (Gadbury et al., 2003).

To observe if the Raman spectra were characteristically different between cohorts, principal component analysis (PCA) followed by linear discriminant analysis (LDA) was used for classification. Our dataset consisted of 1 spectrum/mouse and the dataset was decomposed into principal components (PC) using MATLAB^®^. A standard least-squares fit was applied in JMP®14 to the PCs and the cohort classification to determine what PCs were significantly correlated with cohort classification (Table S1). The significant PCs (1,2,4) were used for a leave one out cross-validation (LOOCV) LDA (Ye et al., 2004) model, using a developed MATLAB^®^ code (Dwinnell, 2010). The classification accuracy was calculated to assess the classification error. For comparison, the classification error using multiple iterations of resampling classifications without replacement was calculated.

To investigate the diagnostic potential of Raman spectroscopy alone or in combination with medical imaging for fracture toughness prediction, we created independent linear regressions using partial least squares regression (PLSR) with a LOOCV approach. We first performed a standard least squares regression in JMP®14 to determine which PCs significantly correlated with fracture toughness (Table S1). LOOCV-PLSR models (Shu et al., 2018), were performed independently three times using, (a) medical imaging data (10×2 matrix), (b) Raman spectral PCs (10×3 matrix), and (c) combined medical imaging and Raman data (10×5 matrix). For simplicity and to guard against over calibration with a small dataset, the model rank was held constant at 2. The data was variance normalized before PLSR. To assess the success of the model, the root mean squared error of cross-validation (RMSECV) and Pearson correlation coefficient (p<0.05) was calculated. For comparison, the RMSECV was calculated from multiple LOOCV-PLSR models with permutated resampled K_c_ values without replacement (Figure S3B-D).

## Results

Structural and compositional properties effects of ovariectomy were determined by μCT, DXA imaging, and Raman spectroscopy. Figure 1 shows that ovariectomy resulted in an 11% decrease in cortical thickness (p<0.05) and a 10% decrease in aBMD (p<0.05), respectively, compared to sham controls. The average Raman spectrum from the OVX cohort (Figure 2A, red) was different from the sham cohort (Figure 2A, blue), as visualized by the spectral difference plot (Figure 2A, green). A closer inspection of the Raman spectral difference plot revealed the sham cohort had a relatively higher signal from phosphate and a relatively lower signal from a peak at 906 cm^−1^. The latter finding is an empirical observation that is still under investigation; such a peak has not been reported in the bone Raman literature to our knowledge.

**Figure 1:**
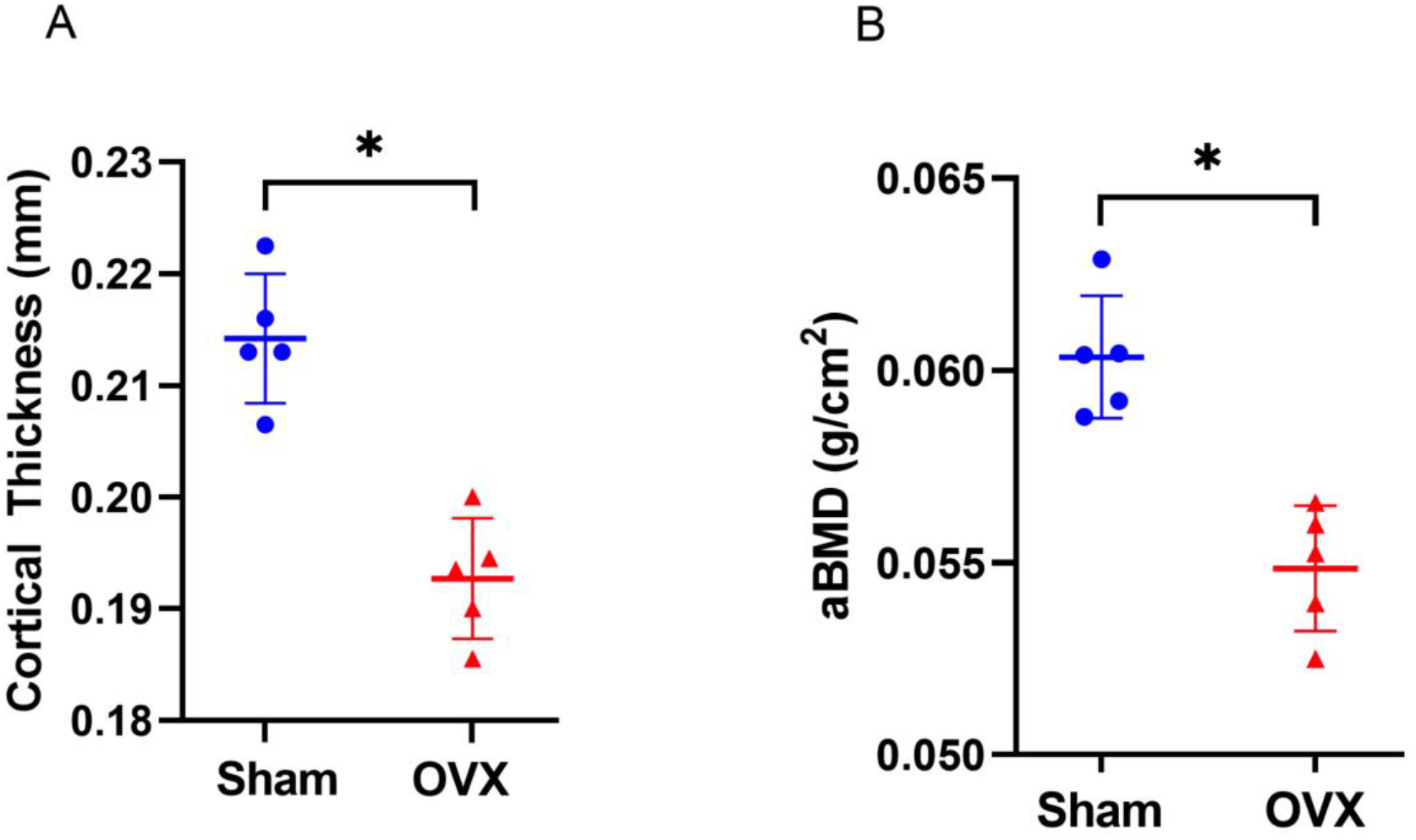
Effects of ovariectomy on femoral bone size and mass. A) Measured cortical thickness from μCT. The OVX cohort had a significantly lower cortical thickness when compared to sham cohort (p <0.05). B) Measured areal bone mineralization density (aBMD) from DXA. The OVX cohort had a significantly lower aBMD when compared to sham cohort (p < 0.05). Data presented as scatter plot due to small sample size (n=5 per cohort) with the mean ± standard deviation. Cohorts compared using permutation tests with significance set at p<0.05.

**Figure 2:**
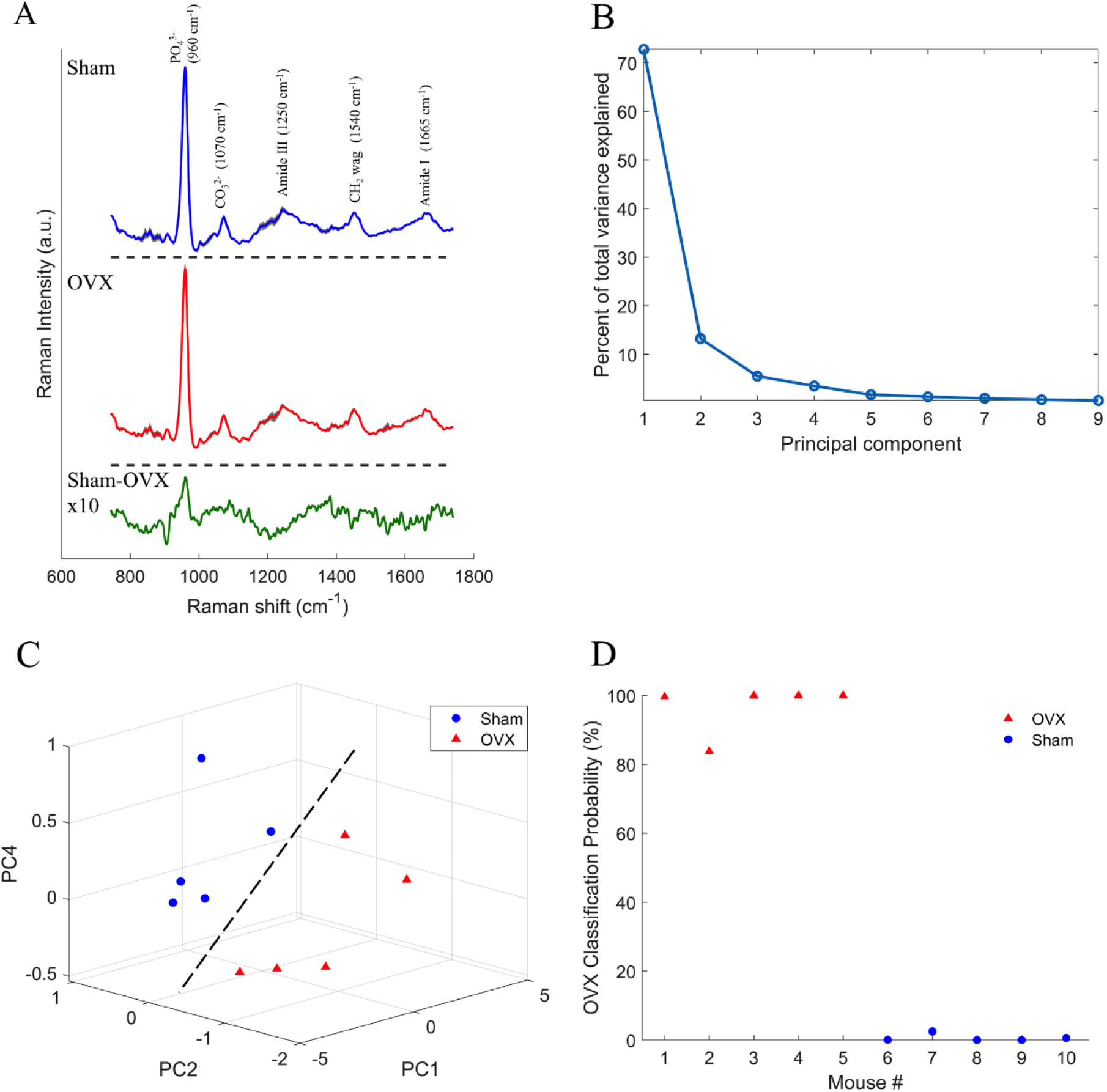
Effects of ovariectomy on femoral Raman spectra and classification analysis. A) Mean sham and OVX spectra (shading represents ± 2×standard deviation) B) Spectral difference plot is magnified by an amplitude of 10 to visualize difference of sham minus OVX mean showing how the mean sham spectrum has a relatively higher phosphate signal when compared to OVX. B) Scree plot showing the percent of total variance explained by each principal component. C) Scatter plot of principal components 1, 2, and 4 showing the samples can be separated based on cohort (dashed line is provided as a guide to the eye; it is not a fit). (D) OVX classification from PCA-LDA using principal components 1, 2, and 4 and LOOCV with 100% accuracy.

Figure 2B shows the principal component contributions to the total variance. Figure 2C shows the 3D scatter plot of PC1, PC2, and PC4 showing a separation based on cohort. The LOOCV-LDA lead to a classification accuracy of 100%, based on surgery (Figure 2D). To determine the stability of our LDA model, we ran multiple iterations with resampling classification without replacement (Figure S3A) showing our LDA results were robust despite the small sample size.

To determine the effects of ovariectomy on fracture toughness, a notch was created on the anterior side of the mid-diaphysis of the femurs (Figure 3A). Three-point bending was then performed to induce a controlled fracture initiating at the experimentally created notch site (Figure 3B). Fracture toughness was computed using measurements of the half-crack angle at fracture instability, average cortical thickness, and mean radii from SEM images (Figure 3C). Consistent with our hypothesis, the fracture toughness constant, *K*_*c*_, was also significantly decreased by 49% (p<0.05) due to ovariectomy (Figure 3D).

**Figure 3:**
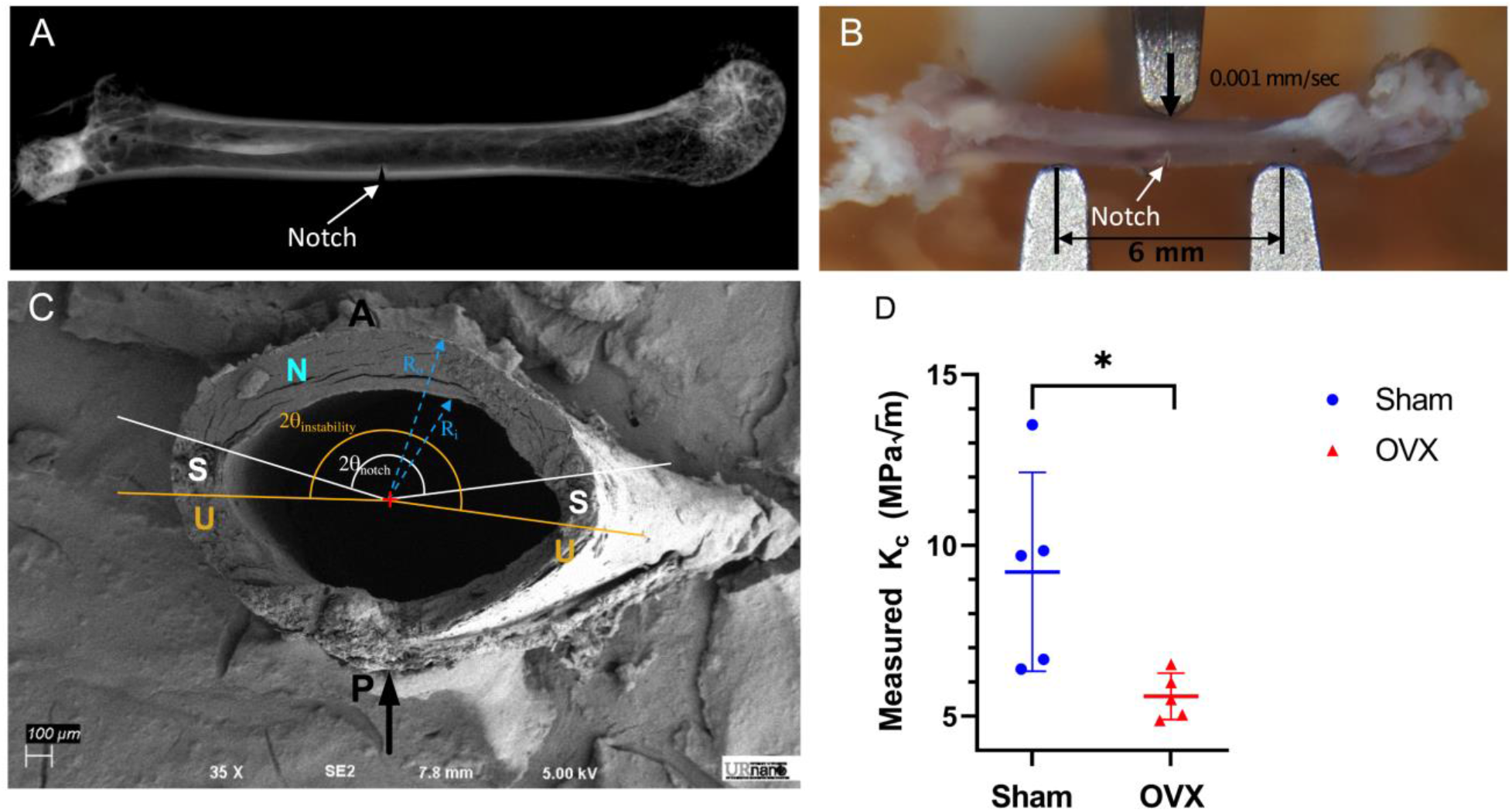
Effects of ovariectomy on fracture toughness. A) X-ray of femur showing notch that was made through the cortical bone in the midshaft. B) Three point bending set-up showing the top post is lined up with the notch to ensure crack propagation occurs at notch. The span length was 6mm and the top plate came down at a rate of 0.001 mm/sec along the posterior-anterior direction. C) SEM image of fracture plane showing regions of stable and unstable fracture propagation. D) Experimentally determined fracture toughness (Kc) showing the OVX cohort had a significantly lower Kc. Data presented as a scatter plot with the mean ± standard error of the mean (n=5 per cohort). Asterisk indicate significant difference determined by permutation test (p < 0.05). Black arrow indicates the direction of loading. P and A denote the posterior and anterior sides of the femur, respectively. N denotes the experimentally induced notch surface. S denotes the stable crack propagation region. U denotes the unstable crack propagation region.

To investigate the improvements afforced by Raman spectroscopy when combined with medical imaging for fracture toughness prediction, we created LOOCV-PLSR models. First, we identified, using standard least squares regression in JMP®14, PCs that significantly correlated with fracture toughness (Table S1). We then showed that the LOOCV-PLSR model to predict fracture toughness using the Raman PCs (PC2,4,5) was significant (p<0.05) with a RMSECV of 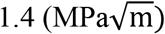 and r^2^ of 0.71 (Figure 4B). Figure 5 shows the PC loadings to demonstrate the parts of the Raman spectra were weighted in the PC analysis. PC2 had distinct peaks for phosphate (mineral) and CH_2_ (matrix), PC4 had peaks for phosphate and carbonate (mineral), and PC5 had a peak for carbonate. In contrast, LOOCV-PLSR models incorporating only aBMD and cortical thickness from medical imaging (DXA and CT, respectively) did not yield a predictive model with a RMSECV of 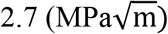 and correlation coefficient r^2^ of 0.16 (p = 0.25) (Figure 4A). Consistent with our hypothesis, we achieved higher accuracy when we combined Raman PCs with aBMD and cortical thickness, the LOOCV-PLSR was a significant model (p < 0.001), with a lower RMSECV of 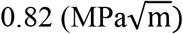 and higher r^2^ of 0.92 (Figure 4C). This improved model predicted that the average fracture toughness of the OVX was 49% lower than the sham mice, which was statistically significant (p < 0.05) (shown in Figure 4D). To determine the stability of our PLSR model, we ran multiple iterations with resampling K_c_ without replacement (Figure S3B-D) showing our reported PLSR results using Raman PCs alone, and the combined model were robust despite the small sample size.

**Figure 4:**
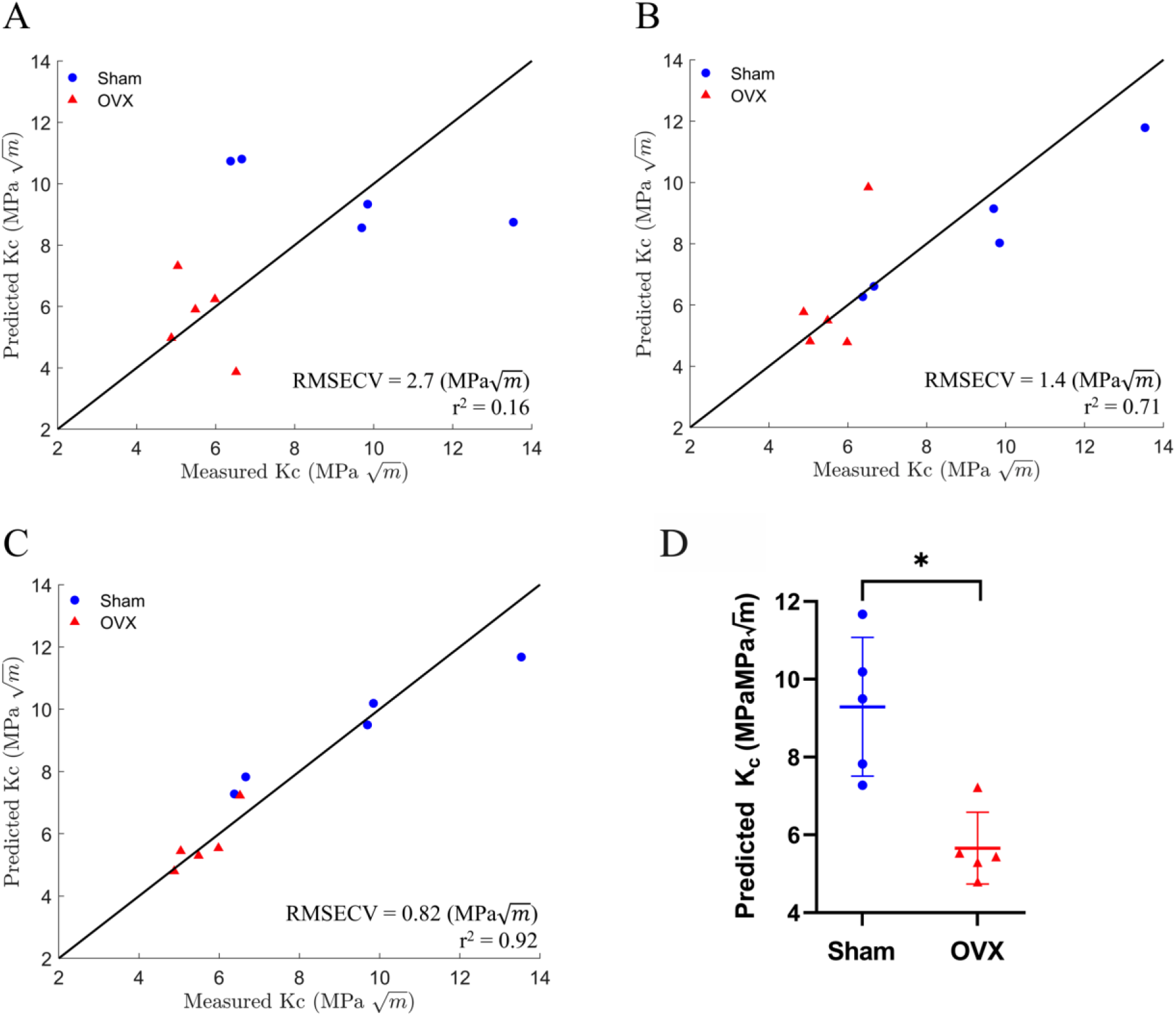
PLSR results for predicting fracture toughness (Kc) with a leave one out cross-validation using: A) BMD and cortical thickness 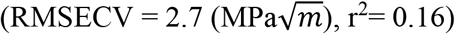 showing no significant correlation (p=0.25). B) Principal components 2, 4, and 5 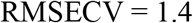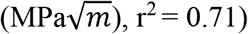 showing significant correlation (p < 0.05). C) Combined PLSR model of aBMD, cortical thickness, and principal components 2,4, and 5 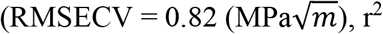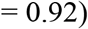 showing significant correlation (p < .0001). Solid line is provided as a guide to the eye; it is not a fit. D) Predicted Kc using model with aBMD, cortical thickness, and principal components. Averaged predicted OVX fracture toughness was significantly lower than sham, determined by permutation test with significance set at p < .0001.

**Figure 5:**
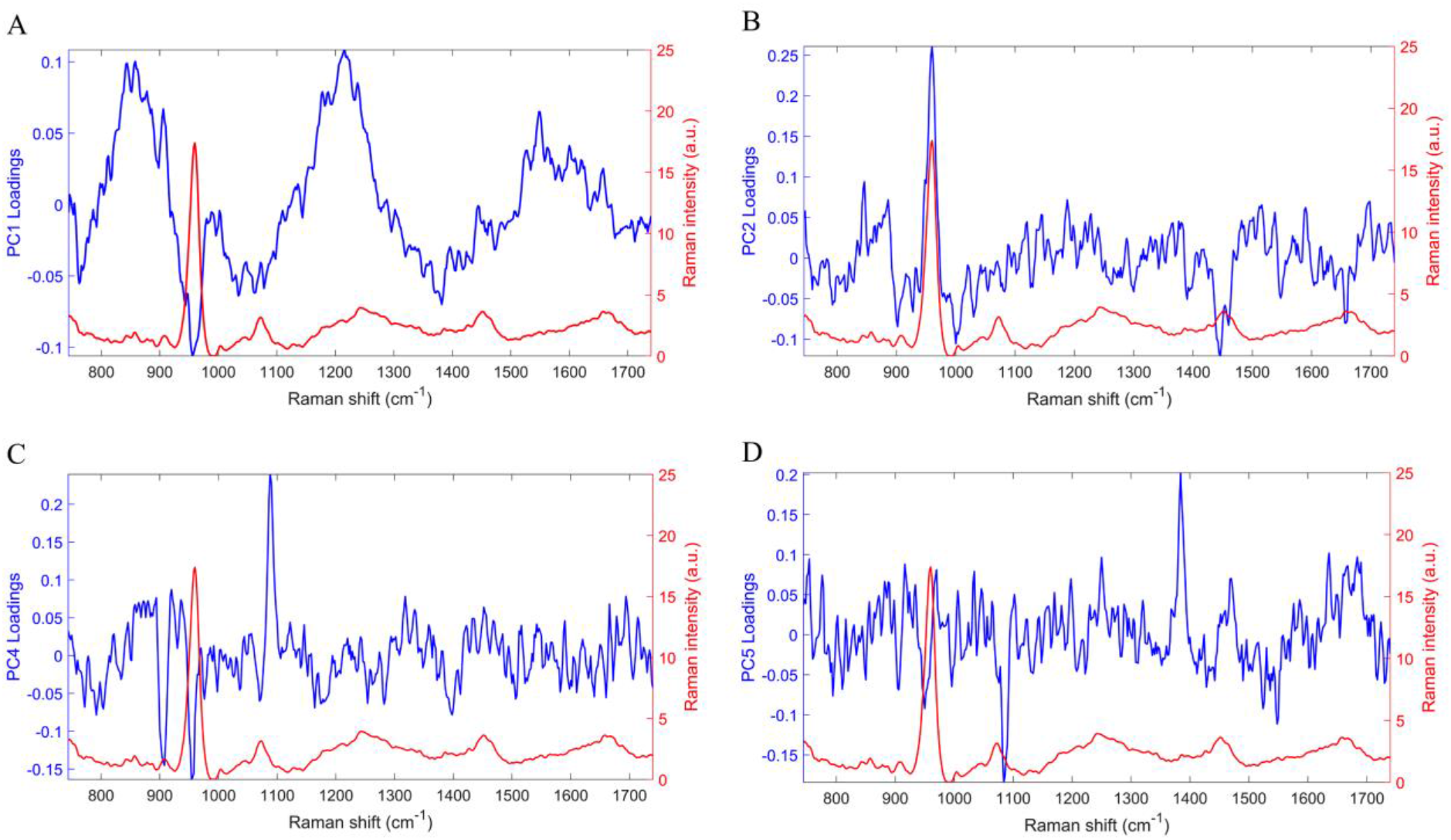
Principal component loadings of PCs that were used for either cohort or Kc prediction (blue) and bone spectrum (red). (A) PC1 was used for cohort classification and shows peaks for phosphate, which contributes to the mineral component of the bone, and amide III which contributes to the matrix component. (B) PC2 was used for both cohort and Kc prediction showing peaks corresponding to phosphate and CH2, which contributes to the matrix component. (C) PC4 used for both cohort and Kc prediction shows peaks for phosphate and carbonate, another mineral component. (D) PC5 used for Kc prediction shows peaks for carbonate.

## Discussion

In this study, we report structural and biochemical changes in femurs due to OVX via DXA, μCT, and Raman spectroscopy. We have shown that OVX was associated with a reduction of femoral aBMD, cortical thickness, and fracture toughness. The Raman spectral based classification accurately discriminated the OVX cohort from the sham cohort with remarkable accuracy (100%). We showed that the bone loss lead to a reduced fracture toughness associated with reduced bone density and cortical thickness. Others have reported OVX decreased tibial cortical thickness (Roberts et al., 2019) and a non-significant reduction of femoral fracture toughness (Shahnazari et al., 2010). Additionally, we show that Raman spectroscopy, especially when combined with DXA and CT, can provide an accurate prediction of fracture toughness deterioration in response to ovariectomy in mice. These results suggest that combining medical imaging with Raman spectroscopy has the potential to improve prediction of cortical bone fracture toughness and potentially the risk of fragility fractures.

We have previously shown that Raman spectroscopy can predict femoral fracture toughness for a rheumatoid arthritis model (Inzana et al., 2013). Others have demonstrated that Raman-derived features can be incorporated in multivariable linear models to predict up to 50% of the variance in human cortical bone’s fracture resistance over a range of ages (Unal et al., 2019). PCs derived from full Raman spectra of human cortical bone specimens have been shown to correlate strongly with fracture mechanics properties such as crack growth toughness and initiation (Makowski et al., 2017). These reports and our findings suggest that Raman spectra should be mined for bone quality assessment.

Despite the small sample size in our study, our robust statistical analysis mitigates this concern. However, the classification accuracy achieved through PCA-LDA of the whole spectra merits further investigation and validation. Future studies should establish this potential and explore more non-linear classification models using machine learning algorithms These studies will need large training data sets and should strongly consider the focus on establishing such classification models based on human bone samples representing a wide spectrum of bone quality. Future studies will include additional measurements of the bone’s matrix composition to understand why Raman helps predict fracture toughness.

Another limitation is that the current study design required the experimental determination of fracture toughness as the ground truth in terms of the ability of the bone to resist fractures, which we loosely associate with fracture risk. While this was necessary for the current cross-sectional study, future studies in mice can be longitudinal. The key to enabling such longitudinal measurements of Raman spectra is the ability to make such measurements through the skin and soft tissue overlying the bone. Our group and others have shown the feasibility of non-contact transcutaneous measurements through the use of spatially offset Raman spectroscopy (SORS) (Feng et al., 2017; Matousek and Stone, 2016; Shu et al., 2018) and novel algorithms to retrieve the bone spectra from the composite spectra such as simultaneous, overconstrained, library-based decomposition (Maher et al., 2013). These advancements can also potentially enable clinical prospective studies of the diagnostic utility of non-contact transcutaneous SORS in classifying patients who might be at higher risk of fragility fractures. A major hurdle to overcome is the ability of the light to penetrate the soft tissue covering the bone in humans, which we believe to be a surmountable hardware design problem.

## Conclusions

In this exploratory study involving ovariectomy-induced osteoporosis in mice, we report LDA Raman-based classification of the bones according to their health. We demonstrate that OVX resulted in a significantly lower fracture toughness and that combining Raman spectroscopy with standard medical imaging outcomes such as aBMD and bone size (cortical thickness) can improve the prediction of fracture toughness, and by corollary help identify fragility fracture risk. To our knowledge, this is the first demonstration of predicting such changes with Raman spectroscopy and medical imaging.

## Conflict of Interest Statement

The Authors have no conflicts to disclose.

## Acknowledgements

The authors would like to thank Ms. Karen Bentley and Mr. Michael Thullen for their technical assistance with SEM imaging and micro CT, respectively. The study was supported by grant numbers R01AR070613, R21AR061285, and P30AR069655 from NIAMS/NIH. The content is solely the responsibility of the authors and does not necessarily represent the official views of the NIH.

## Notes

### Competing Interest Statement

The authors have declared no competing interest.

