## Supplementary Material for "Improved prediction of femoral fracture toughness in mice by combining standard medical imaging with Raman spectroscopy"

### Supplemental Data

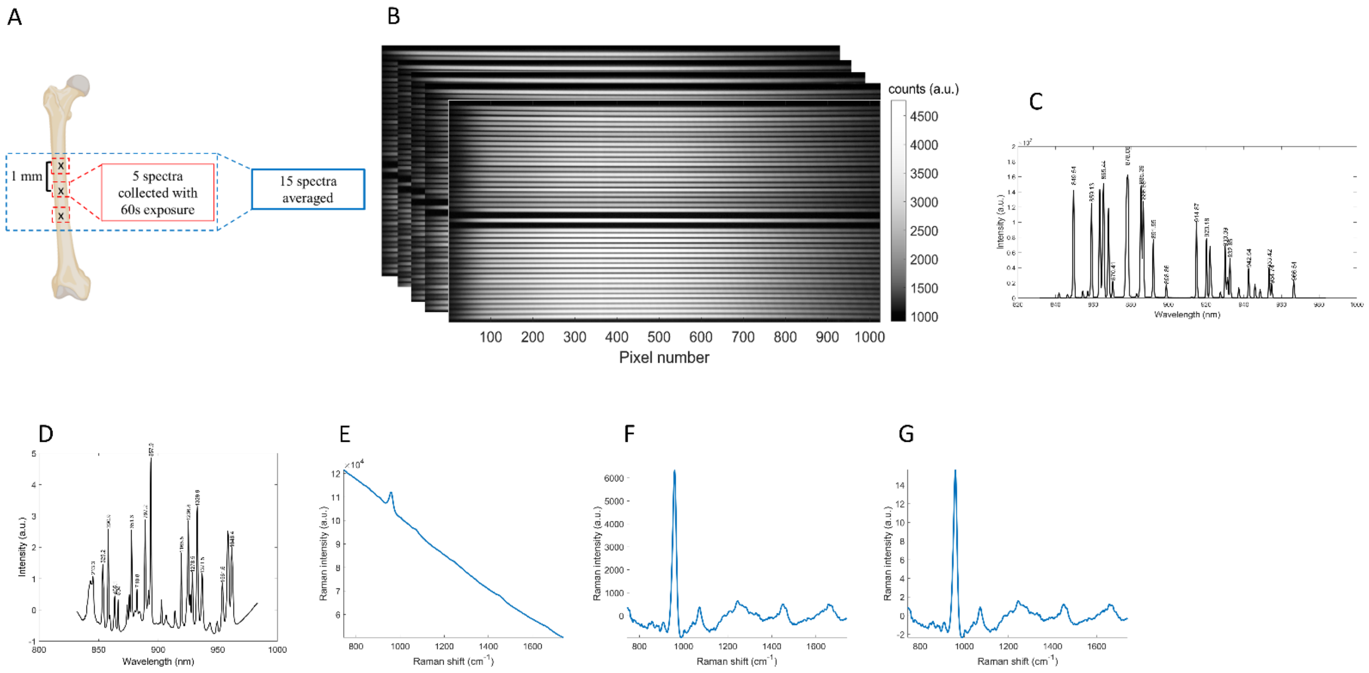


**Figure S1:** Raman data acquisition and spectral pre-processing procedure

Figure A shows 3 locations were measured along the anterior side of the femur spaced 1 mm apart. Each location consisted of 5 spectra acquired with 60 second exposure. Figure B shows the images collected from our CCD spectrograph. The five images correspond to each 60- second exposure. Each horizontal stripe represents a spectrum collected from an optical fiber. To correct for readout and dark currents that are present during data acquisition from the CCD, a dark image is acquired separately with the laser off. This image is subtracted from measured CCD images.

To calibrate for wavelength, a neon, gas-discharge lamp (Model 6032, Newport Corporation, Irvine, CA). A third-polynomial regression was used to perform a calibration from pixel to wavelength shown in figure C.

To determine the Raman shift, TYLENOL® (McNEIL-PPC Inc., Fort Washington, PA) was measured (figure D). We used the relationship between Raman shift (Δν), spectral wavelength (λ), and wavelength of the illumination source ($\lambda_0$) shown below:

$$\Delta\nu = \frac{1}{\lambda_0} - \frac{1}{\lambda}$$

Correction for aberrations in the spectrograph was conducted by using a method by Esmonde-White et al. (Esmonde-White et al., 2011). A metal-ion-doped glass (Standard Reference Material 2241, National Institute of Standards) was used for scaling purposes. For the glass spectral image, each row was divided by the known emission spectrum determined by the

National Institute of Standards. Intensities of the vertical strip of pixels were fit with 40 Gaussian-Lorentzian line shapes (one for each fiber) to determine the spectral throughput associated with each fiber. Raman spectra were extracted from spectral images using a method described by Dooley et al (Dooley et al., 2010). Specifically, the Gaussian-Lorentzian line shapes from the glass image were fit to the measured CCD image using a least-squares fit. The uncertainty in each fit was calculated using a previously described method (Scepanovic et al., 2007) and the spectrum was scaled such that the square root of the mean spectral intensity was equal to the ratio of mean spectral intensity to the fit-uncertainty. Figure E shows a raw Raman spectrum (744-1740 $cm^{-1}$), spectra were smoothed with a Savitzky-Golay filter (Savitzky and Golay, 1964).

The fluorescence is then fit by a $5^{th}$ order polynomial and subtracted off shown in Figure F. Spectra were normalized to the mean absolute deviation (MAD) with respect to their mean, determined by:

$$MAD = \frac{1}{n}\sum_{i=1}^{n} | S_i - \bar{S} |$$

was equal to 1, where S is the spectral intensity shown in figure F.

**<u>References</u>**


1. Dooley, K., Esmonde-White, F.W., Morris, M., 2010. Optical fiber bundle coupling errors in Raman spectra: correction via data processing. SPIE.
2. Esmonde-White, F.W., Esmonde-White, K.A., Morris, M.D., 2011. Minor distortions with major consequences: correcting distortions in imaging spectrographs. Appl Spectrosc 65, 85-98.
3. Savitzky, A., Golay, M.J.E., 1964. Smoothing and Differentiation of Data by Simplified Least Squares Procedures. Analytical Chemistry 36, 1627-1639.
4. Scepanovic, O., Bechtel, K., Haka, A., Shih, W.-C., Koo, T.-W., Berger, A., Feld, M., 2007. Determination of uncertainty in parameters extracted from single spectroscopic measurements. Journal of biomedical optics 12, 064012.

**Table S1:** p values from standard least squares regression results

| | Significance for cohort class | Significance for $K_c$ |
|---|---|---|
| PC1 | **0.03225** | 0.06419 |
| PC2 | **0.02986** | **0.03335** |
| PC3 | 0.28943 | 0.09695 |
| PC4 | 0**.03295** | **0.03140** |
| PC5 | 0.09004 | **0.03793** |
| PC6 | 0.07905 | 0.16051 |
| PC7 | 0.087227 | 0.13493 |
| PC8 | 0.07084 | 0.18226 |

Standard least squares regression (no cross validation) in JMP® 14 to determine a combination of PCs that were correlated with cohort and $K_c$. The p-value given tests the hypothesis that there is no relationship between the predictor (PC) and the response (cohort or $K_c$). With a significance level set at 0.05, we determined PC1, PC2, and PC4 were significant predictors of cohort. A scatter plot of the three PCs showed the cohorts could be separated, and a LOOCV-LDA model yielded a 100% classification accuracy. PC2, PC4, and PC5 were significant predictors of $K_c$ and this was validated with our LOOCV-PLSR model.

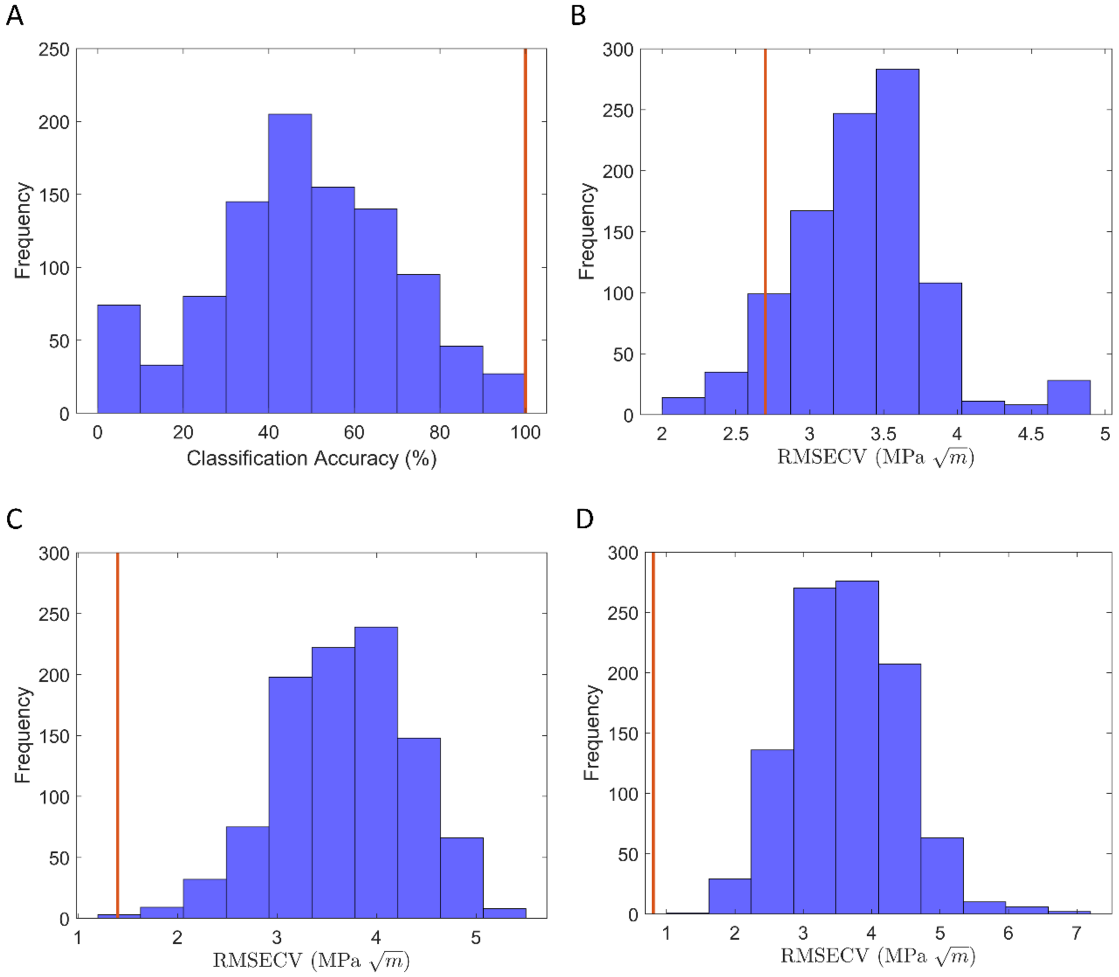


**Figure S2**: Classification (A) and regression (B-D) results from permutation. Samples' cohort class and $K_c$ values were resampled without replacement 1000 times and the LOOCV-LDA and PLSR models were run each time to calculate classification accuracy and RMSECV respectively. Histograms were then generated to show the distribution of prediction results, with the red line denoting where our reported results fall within the distribution. For our actual classification results using LDA, the 100% accuracy falls within the 99.5$^{th}$ percentile of the permutation distribution (A), supporting the claims made in the manuscript that our results are robust despite the small sample size. For Kc prediction using aBMD and cortical thickness alone, the actual RMSECV of 2.7 (MPa$\sqrt{m}$) falls within the 75$^{th}$ percentile of the permutation distribution. The manuscript reports our model using X-ray yielded no significant correlation RMSECV = 2.7 MPa$\sqrt{m}$) and $r^2$= 0.16 (p = 0.16). Permutations using 3 PCs alone place the actual RMSECV of 1.4 (MPa$\sqrt{m}$) within the 99$^{th}$ percentile of the distribution (C). The manuscript reports our model using Raman PCs as significant ($p<0.05$) RMSECV = 1.4 MPa$\sqrt{m}$) and $r^2$= 0.16 (p = 0.71). Permutations on the combined model of 3 PCs, aBMD, and cortical thickness place the actual RMSECV of 0.81 (MPa$\sqrt{m}$) within the 99$^{th}$ percentile of the distribution (D). The manuscript reports the model using both Raman and X-ray as significant ($p<.0001$) RMSECV = 0.82 MPa$\sqrt{m}$) and $r^2$= 0.92. These results support the claims made in the manuscript that our results are robust despite the small sample size.
